# BioConvert: a comprehensive format converter for life sciences

**DOI:** 10.1101/2023.03.13.532455

**Authors:** Hugo Caro, Sulyvan Dollin, Anne Biton, Bryan Brancotte, Dimitri Desvillechabrol, Yoann Dufresne, Blaise Li, Etienne Kornobis, Frédéric Lemoine, Nicolas Maillet, Amandine Perrin, Nicolas Traut, Bertrand Néron, Thomas Cokelaer

**Affiliations:** Institut Pasteur, Université Paris Cité, Plate-forme Technologique Biomics, F-75015 Paris, France; Institut Pasteur, Université Paris Cité, Bioinformatics and Biostatistics Hub, F-75015 Paris, France; Institut Pasteur, Université Paris Cité, G5 Sequence Bioinformatics, Paris, France; Institut Pasteur, Université Paris Cité, G5 Evolutionary Genomics of RNA Viruses, Paris, France; Institut Pasteur, Université Paris Cité, CNRS UMR3525, Microbial Evolutionary Genomics, Paris, France; Institut Pasteur, Université Paris Cité, Unité de Neuroanatomie Appliquée et Théorique, F-75015 Paris, France

## Abstract

Bioinformatics is a field known for the numerous standards and formats that have been developed over the years. This plethora of formats, sometimes complementary, and often redundant, poses many challenges to bioinformatics data analysts. They constantly need to find the best tool to convert their data into the suitable format, which is often a complex, technical and time consuming task. Moreover, these small yet important tasks are often difficult to make reproducible. To overcome these difficulties, we initiated **BioConvert**, a collaborative project to facilitate the conversion of life science data from one format to another. **BioConvert** aggregates existing software within a single framework and complemented them with original code when needed. It provides a common interface to make the user experience more streamlined instead of having to learn tens of them. Currently, **BioConvert** supports about 50 formats and 100 direct conversions in areas such as alignment, sequencing, phylogeny, and variant calling. In addition to being useful for end-users, **BioConvert** can also be utilized by developers as a universal benchmarking framework for evaluating and comparing numerous conversion tools. Additionally, we provide a web server implementing an online user-friendly interface to **BioConvert**, hence allowing direct use for the community.

## 1 Introduction

Over the past few decades, bioinformatics has produced a large number of open source software tools and formats within the academic community. Twenty years ago, in 2002, [1] was already making this observation. Nowadays, these tools, including those designed specifically for Next Generation Sequencing (NGS), are available in various flavors and levels of complexity: single software [2], libraries for developers [3], suite of applications on websites [4], and set of NGS workflows [5, 6]. Installing and using these tools can be challenging in many ways. In particular, it requires knowledge of numerous biological formats, making it very challenging for life scientists to i) make the files suitable for the input of a given tool, ii) connecting the output of one tool to the input of an other tool, iii) to extract metadata from output files for subsequent analysis. To address these challenges, the open source community has made significant progress in different areas. First of all, in the software implementation and installation perspective, great efforts have been made to provide reusable and documented software. For example, the Bioconda [7] project offers a large selection of bioinformatics packages (over 7,000) that are pre-compiled for various platforms. Other resources, such as Python’s Pypi, R’s Bioconductor, and Perl’s CRAN, also provide a wealth of packages for common programming languages in life sciences. Additionnaly, large efforts have been made in the inventory, classification and description of bioinformatics tools. For example, EDAM ontology [8] has been developed to classify and describe bioinformatics tools and data formats in a machine understandable way. More recently, Bio.tools [9] has been designed to provide a catalogue of bioinformatics tools with their annotations (including EDAM), in order to facilitate the search for tools to be integrated in workflows. Despite the abundance of resources, it can still be difficult for new users to navigate this dynamic environment and identify the best software tools for a given task. Even experienced users may face challenges with the continuous emergence of new software and the potential for older tools to become obsolete or lack maintenance. In particular, complex NGS analysis often requires the use of many conversion tools, increasing the knowledge and expertise required to perform conversions across various omics data types (e.g., genomics, transcriptomics, epigenetics).

Web services are frequently used in the field of life sciences, with the NCBI and EBI institutes being two significant providers of online tools and services, including many conversion tools. For example, the EBI’s *seqret* service [10] allows users to convert various standard sequencing formats, including those that are no longer supported. While users can typically only upload a small number of files at a time, these online resources can also be accessed through programming. For instance, the BioServices library [11] allows Python users to programmatically access web services from within their code. Although useful, relying on online resources may not be the best solution as compared to standalone applications. Indeed, the application programming interface (API) may change over time and may also suffer from delay due to the upload and download of large files (a common situation in life sciences). Therefore, local standalone software is usually a far more efficient solution.

Another challenge for users is that new data formats are frequently introduced as new technologies emerge (for example, the FAST5 format from Nanopore [12], the consensus file from PacBio [13]). A community centered around a common software tool for converting life sciences data would enable the integration of new technologies and software at a faster pace. There are already many conversion tools currently available, such as ReadSeq [14], SBFC [15], Seqret, Gotree/Goalign [16], etc. However, they are dispersed, each of them only support a limited number of formats, they may be difficult to install, or may not be optimized for efficient use (*e*.*g*., to be applied on huge datasets).

To overcome these limitations, we initiated **BioConvert**, a collaborative project that aims at providing a common interface for converting life science data from one format to another. It currently supports 50 formats and 100 conversions across various expert knowledge domains such as variant calling, phylogenetics, sequencing and sequence alignments. In this manuscript, we describe the methodology we employed and design choices that we have implemented to facilitate the easy integration of new format conversions into **BioConvert**. We also demonstrate the ability to benchmark multiple methods for specific conversions, in order to select the most efficient tool for each format conversion.

## 2 Materials and Methods

**BioConvert** is a software library written in Python that utilizes an object-oriented approach, where all conversions inherit from a common class. This design ensures that developers follow a consistent set of guidelines and makes it easier to extend the library over the long term. Our primary goal while designing **BioConvert** was to provide a single command line interface for end-users, as demonstrated in the Results section.

**BioConvert** provides two main functionalities that are described below: i) a format conversion framework designed to convert data between formats, and ii) a benchmark framework designed to test the efficiency of all integrated tools and help selecting the most efficient tools for each conversion.

### 2.1 Conversion framework

Regarding the conversion framework, we provide two kinds of implementations. The first ones consists of original implementations developed in Python within **BioConvert**, and are well adapted for simple conversions such as FastQ to FastA sequencing data format. These native implementations have the benefit of reducing the number of external dependencies. The second kind of implementations transparently integrates external tools in **BioConvert**’s own structure and call them by interacting with the host system. This approach is well adapted to more advanced conversions involving more complex formats. In this case, it is often more efficient to use external tools, especially when they are well-established and known to be effective. For example, **BioConvert** uses SAMtools [3] or BAMtools [17] software to perform conversions of read alignment formats (SAM and BAM). This approach allows **BioConvert** to leverage the strengths of existing tools while still providing a unified interface for users, which makes it highly suitable for integration in large workflows (see Supplementary).

Since **BioConvert** relies on external libraries, changes to those libraries may affect the conversion process. To mitigate this risk, **BioConvert** includes a comprehensive set of tests, together with test data, that can be easily extended. This is a common practice for large libraries, and **BioConvert** encourages its usage. To perform these tests, input and output files for each converter are required. These files are used for the testing suite, but they can also act as examples for users. They are typically kept small to facilitate automatic testing with a continuous integration running on a weekly basis or as soon as a change occurred in the library.

**BioConvert** was designed for easy integration of new conversions. Technically, each conversion derives from a single class that defines common methods and attributes. As shown in Listing 1, a conversion from format A to B is named A2B (line 3) and must have at least one valid method performing the conversion. However, multiple methods can be defined, typically for providing several alternative tools (see lines 7 and 11). Methods must be named with the `_method` prefix followed by a user-defined identifier, and they can either use external libraries or be implemented in pure Python. As shown in listing 1, it is possible to write a new conversion in just a few lines of code.

**Listing 1:**
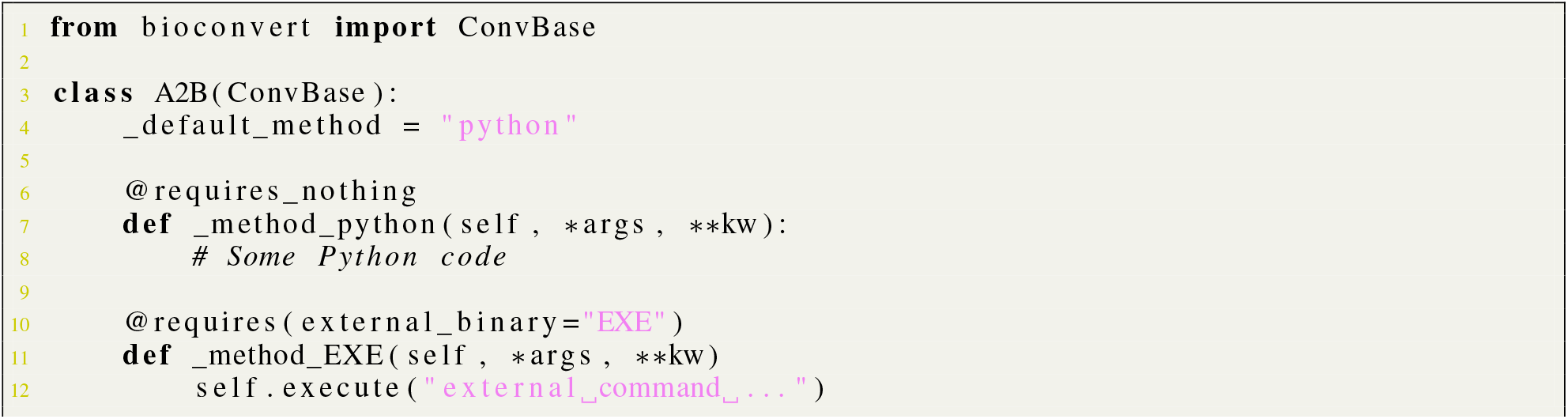
Template of a new converter performing conversion from format A to format B. Methods are implemented using Python and optionally external binaries.

In **BioConvert**, we have implemented a framework that automatically analyses code to detect new conversions, making it easy for developers to add their own methods. When integrated in the **BioConvert** library, the code is automatically parsed to extract the necessary information about the converter: i) the name of the conversion (in this case, A2B), ii) its input and output formats (A and B), and iii) the default conversion method. This information is then made available for users of the `bioconvert` standalone (*e*.*g*., in the command help), with no additional work required by the developers. In the example given in Listing 1, we present a conversion from format A to B, and show how to implement two methods called “python” and “EXE”. It is worth noting that **BioConvert** allows the integration of an unlimited number of methods for a given conversion. That is why indicating a default method is important for typical end-users (in Listing 1, the “python” method is defined as the default on line 4). One of the major advantages of having multiple methods available for a single conversion is that developers can easily benchmark their methods against those already available in **BioConvert** using our conversion benchmark framework.

### 2.2 Benchmarking framework

Benchmarking is a key aspect of **BioConvert**, that can be performed in several ways: from within the library (for developers), from the **BioConvert** command line interface (for end-users), or through a Snakemake [18] pipeline for more intensive testing. For a given conversion and method, benchmarking results can be impacted by various factors, including the size of the input file, the type of data, the number of threads, and the CPU and hard drive performances. Some external tools may also require initialization time before processing the data. To help developers compare their methods to those already available in **BioConvert** in the fairest way, we have implemented a benchmarking procedure that runs conversions multiple times and computes distributions (see Results for more information).

To facilitate better comparisons, we also provide some larger input files for benchmarking purposes. Output files are not required, as the focus of this specific test is not on measuring the exactitude of the conversion, but rather on measuring the computational time required to process input files. Since input files may be large and multiple conversions may be performed, it is not practical to store these files within the **BioConvert** library. To easily access these files, we have created a Zenodo community https://zenodo.org/communities/bioconvert/ to publicly store benchmarking files. These files are only provided when more than one method has been implemented for a given conversion.

## 3 Results

### 3.1 An extensible and robust Python library

**BioConvert** was developed using the Python programming language to create a flexible and extensible library. The object-oriented approach of the language was used to implement a common parent class shared by all converters (see Listing 1). The library provides extensive documentation available online at https://bioconvert.readthedocs.io, updated automatically after each modification of the library. It is important to ensure that conversions remain valid after any update. Therefore, we have included a large set of tests that are run whenever the library is updated. The coverage rate of this test suite is higher than 90%, which is a high standard. It is integrated with the GitHub CI action framework, with one workflow per conversion, making the integration process fast and modular.

### 3.2 An intuitive command line tool

**BioConvert** is designed to be simple and intuitive for end-users, with the goal of providing a common syntax for all conversions that requires as few arguments as possible. To achieve this, we have implemented an implicit mode where the type of conversion is inferred from the input and output file extensions. For example, the following command converts an input FastQ file to an output FastA file, without explicitly specifying the type of conversion:

**Listing 2:**
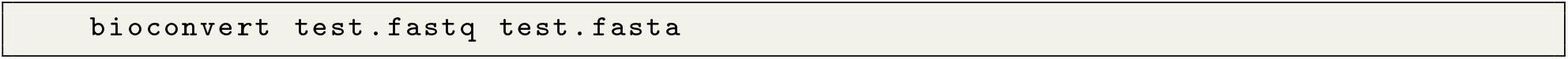
Example of an implicit conversion where extensions suffice for inferring the type of conversion required.

The mechanism behind implicit conversions is based on a registry of common file format extensions used in **BioConvert**. Based on the extensions of the input and output file names, **BioConvert** infers the desired conversion. Although it is common for FastQ files to have extensions such as .*fq* and for FastA files to have .*fa* extensions, there may be some variations (.*fastq*, .*fasta*, etc.). This is why it is possible to register multiple extensions for a given format. In cases where users use non-standard extensions, the implicit mode may not be able to resolve the extensions. In these cases, users can switch to the explicit mode by specifying the required conversion manually:

**Listing 3:**
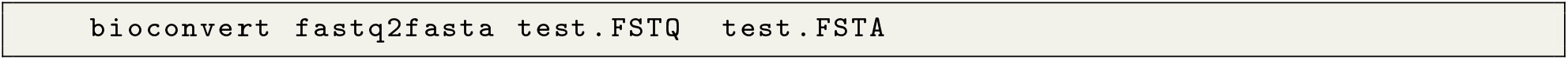
Example of an explicit conversion where extensions can not be resolved automatically.

Because the explicit mode specifies the type of conversion, the second argument can be omitted. Consider for instance the conversion of the SAM alignment format into its binary version (BAM). You can use the explicit mode:

**Listing 4:**
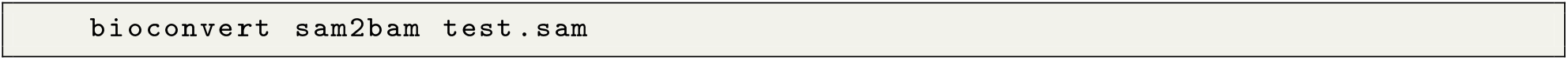
Example of an explicit conversion with an implicit output.

that will create a BAM output file with the same base filename (in Listing 4: “test”) and adding the relevant extension. Of course, you can be explicit again if you need a different output file:

**Listing 5:**
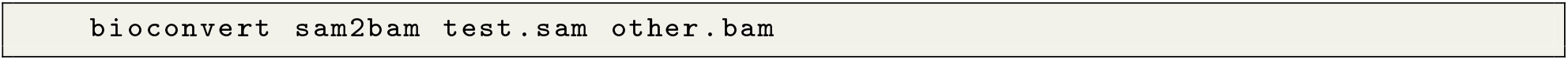
Example of an explicit conversion with an explicit output.

### 3.3 A versatile set of conversions for life sciences

We have described two examples of conversions involving common formats, such as sequencing and read alignment formats such as FastQ, FastA, SAM, and BAM formats. As of the time of writing, **BioConvert**’s latests release includes 50 formats and 100 conversions. Fig. 1 shows a directed acyclic graph representing all possible conversions. Some formats are highly connected due to their common usage in sequence analysis and NGS (e.g., FastQ and FastA). These formats have also served as a starting point for new users to add conversions to the **BioConvert** project. Fig. 2 presents the same graph, with formats and conversions grouped by scientific domains. We identified eight main domains: variant calling, sequencing, phylogeny, assembly, read alignment, coverage, annotation, and compression. Most of the formats in **BioConvert** are related to NGS, due to the projects’s developers specializating in this area. However, formats related to other omics disciplines such as proteomics could also be implemented in **BioConvert**. In Fig. 1, some nodes do not have names (just a small circle). They correspond to logical gates that accept multiple inputs or outputs. For example, the FastQ format consists of both a sequence and its quality. The sequence is stored in FastA format, while the quality information is usually discarded if only the sequence is of interest. In **BioConvert**, it is possible to convert a FastQ file into either its FastA or quality file, or to save both using the following syntax:

**Listing 6:**
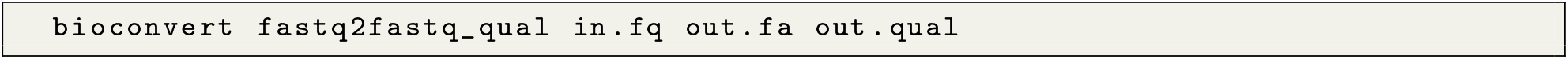
Example of conversion that produces two output files.

**Figure 1:**
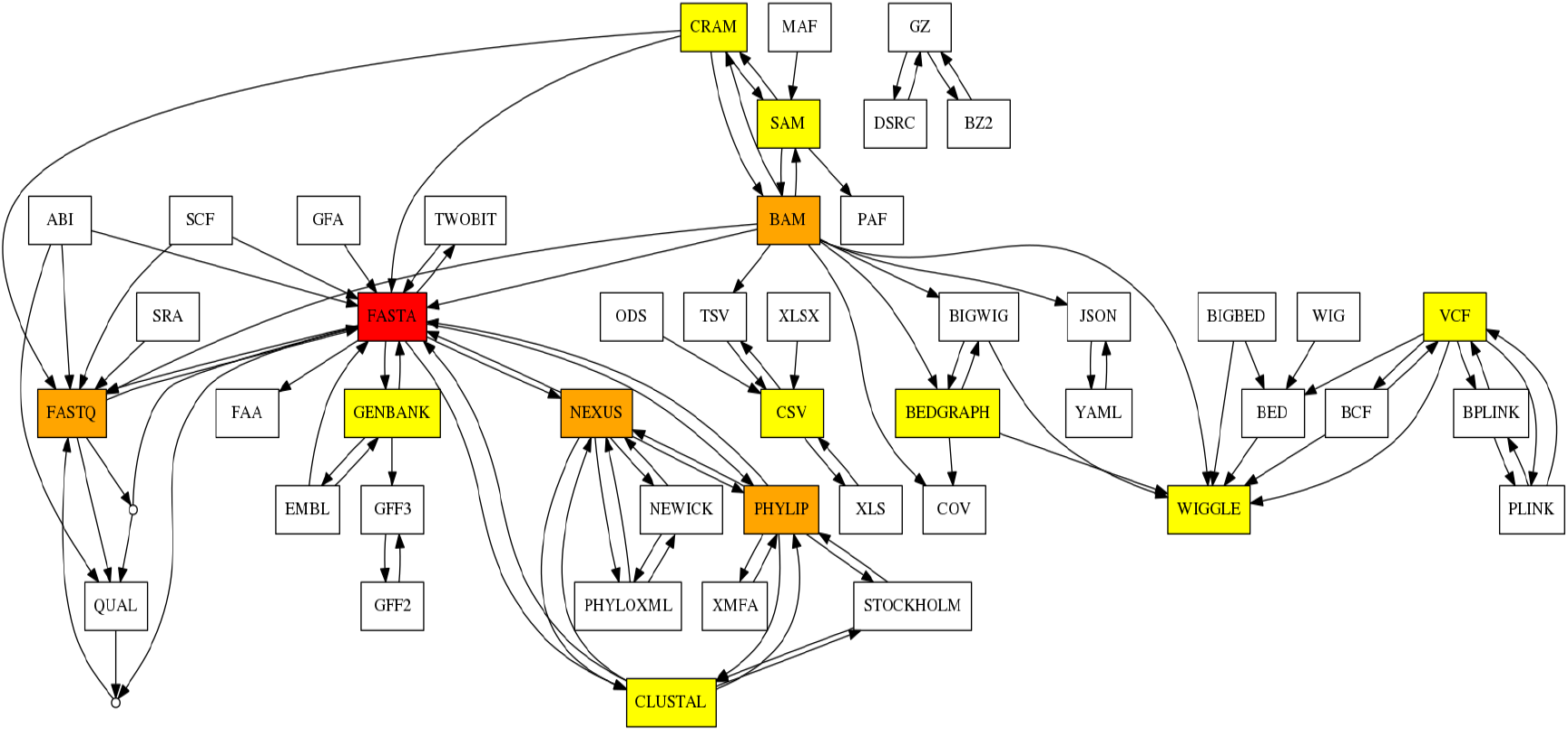
Formats and conversions available in **BioConvert** are represented as a directed acyclic graph. Nodes correspond to formats and edges correspond to conversions. Colors indicate the degree of each format (number of connections/conversions that a node/format has in the graph).

**Figure 2:**
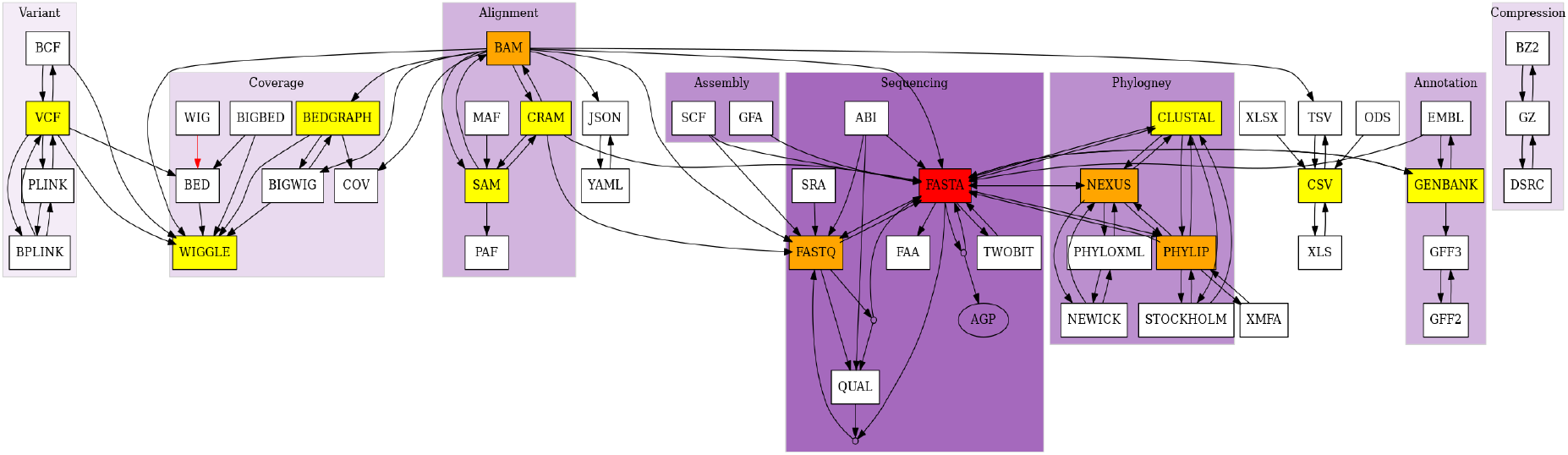
The formats included in **BioConvert** cover NGS formats. In this graph, nodes (formats) are clustered according to their field of expertise. We could identify several topics including variant calling, phylogeny, sequencing data, alignments, …

**BioConvert** also supports compressed and uncompressed input and output files for certain formats. For instance, the `fastq2fasta` conversion requires no extra input from the user. It can detect common compressed file extensions (such as bzip and gunzip), and automatically decompress input files and compress output files based on the given extension.

### 3.4 Benchmarks

In **BioConvert**, most converters have only one method available. However, around 30% of the conversions have 2 or 3 methods, as shown in Fig. 3. One conversion, `fastq2fasta`, has 8 methods, which was used as a practical example during the initial setup of the **BioConvert** framework.

**Figure 3:**
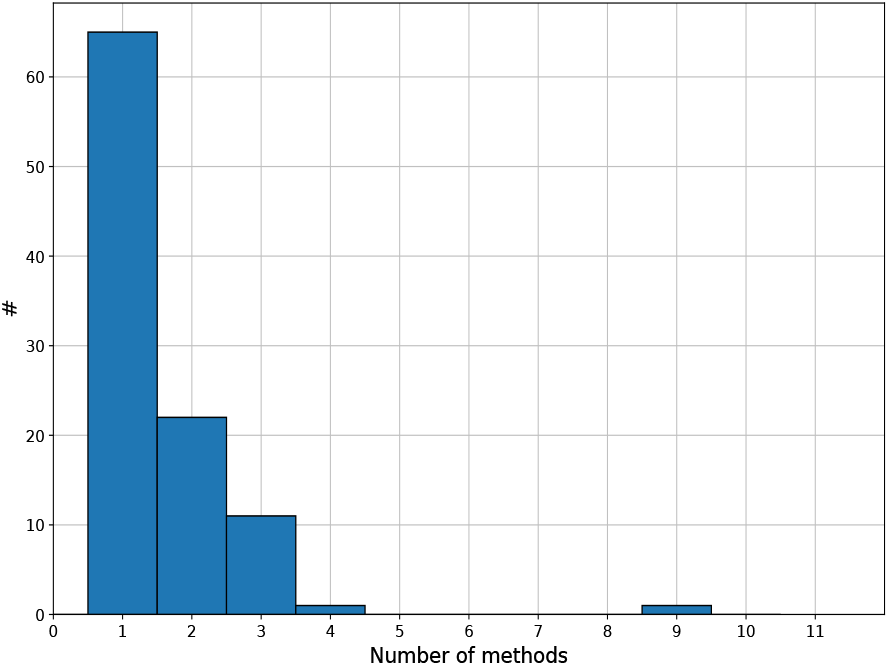
Number of methods implemented in each conversion. Most have only 1 or 2 methods.

When a conversion provides multiple methods, a default method needs to be specified. This is usually based on expert knowledge or the best-performing method. While many benchmarks have been published for specific software tools [19, 20, 21], local computing resources, the input file’s nature, and hard disk performance can all impact benchmarking results. To address this, we have implemented a benchmark framework within **BioConvert** that can be run locally on any input file. This is known as single-mode benchmarking. For example, the benchmark of the `fastq2fasta` conversion (Fig. 4) consists of running each method *N* times (5 by default) to account for fluctuations and calculate the average and standard deviation across all methods. This allows users to determine the best method to use on their own infrastructure. In listings 2-5, users simply need to add an argument (-b) to obtain plots such as the one shown in Fig. 4.

**Figure 4:**
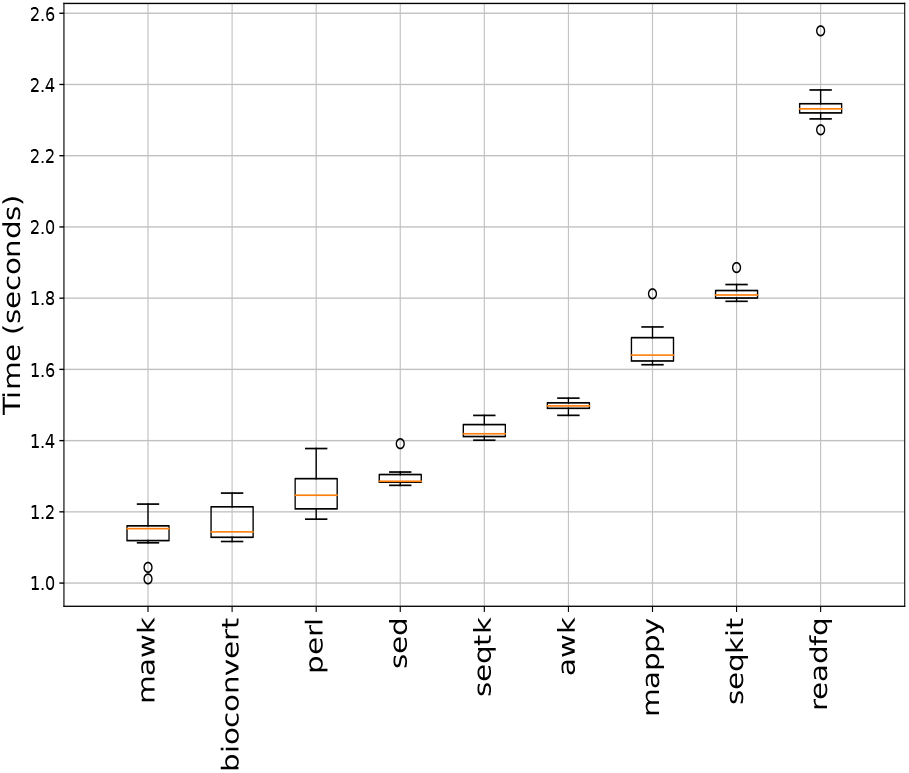
Single-mode benchmarking. **BioConvert** provides a sub-command to compare the computational time of all methods available within a given converter (here FastQ to FastA). Each method is run several times to estimate the average time for each method as well as the standard errors. In this instance, the `mawk` method gives the best performance. Results and error bars may fluctuate depending on hardware performances and concurrent running processes. Benchmark obtained with a SSD hard disk, with compressed input and uncompressed output files.

While the single-mode benchmark is generally adequate, the error bars might be wider than the gap between two groups, and various external factors such as concurrent processes running on the same system can impact the runtime of a particular method in the conversion process. To mitigate this issue and ensure the reliability of the benchmark results, **BioConvert** also implements a multi-mode benchmark that repeats the single-mode benchmark multiple times. This is achieved using a Snakemake [18] pipeline. By doing so, we obtain a more robust estimate of the median run time of each conversion method. The input files (See Benchmarking framework section) used in this benchmark/pipeline are retrieved automatically to ensure that all developers are using the same files. The benchmark can also be run on high-performance computers or locally, with the ability to distribute processes more randomly. In the case of the `fastq2fasta` conversion, we ran the multi-benchmark on a high-performance computing (HPC) infrastructure. As shown in Fig. 5, some methods had fluctuations when run multiple times, but others were more stable, such as the mawk method which showed low variability of the median run time.

**Figure 5:**
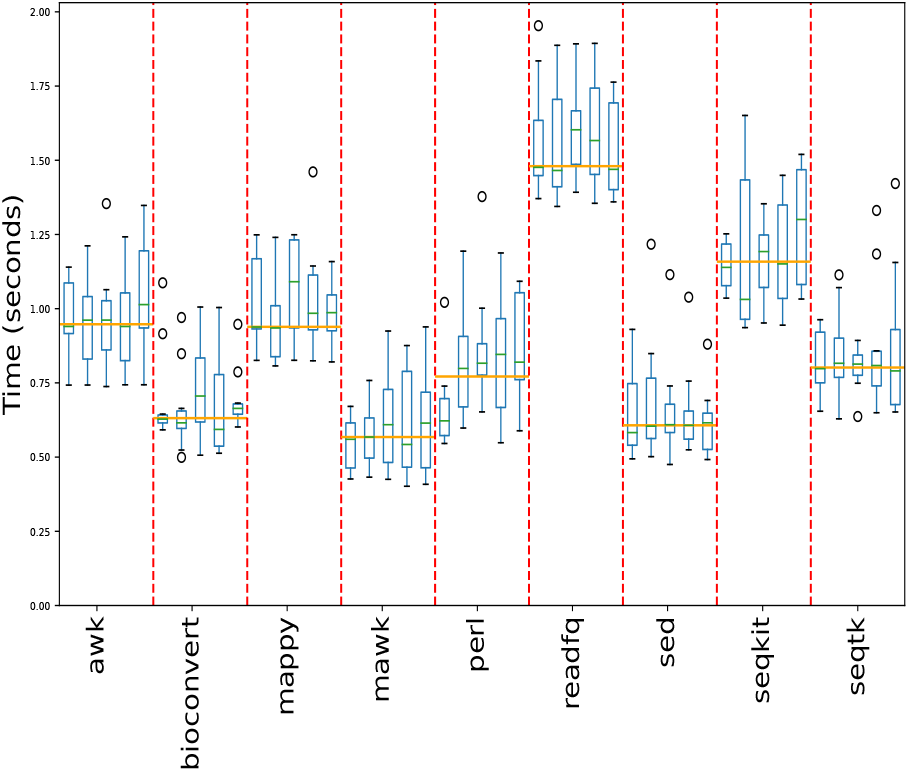
The multi-mode benchmark of the `fastq2fasta` conversion involves repeating the single-mode benchmark multiple times to better understand the variability within a method and provide more confidence in determining the fastest method. The mawk method has consistently low variation and is one of the fastest methods.

The benchmark framework in **BioConvert** is useful not only for comparing the performance of different conversion methods but also for optimizing the local installation of **BioConvert**. Developers can use the framework to verify previous benchmark results or to evaluate the impact of software updates. For instance, in Fig. 6, the conversion from BAM to SAM format was benchmarked using a local installation of samtools version 1.7. The results showed that `sambamba` was set as the default. However, when `samtools` was updated to version v1.15, the benchmark results changed significantly, and the default method was updated accordingly.

**Figure 6:**
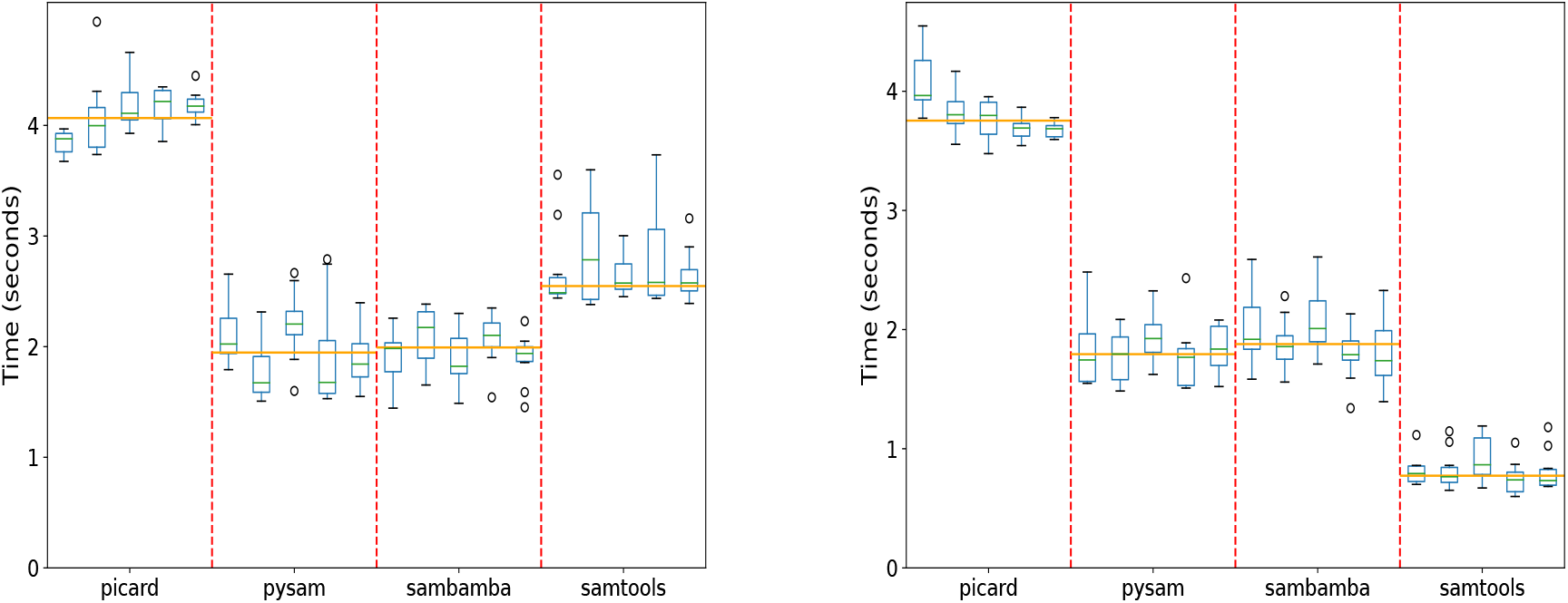
Benchmarking of the `bam2sam` converter with four methods implemented in **BioConvert**. The only difference between top and bottom panels is related to the version of samtools used for benchmark (1.7 and 1.15 respectively). In the top panel, the performance of `samtools` and `sambamba` were similar, while in the bottom panel, `samtools` was 2-3 times faster. This difference can be attributed to the updated version of `samtools` resulting in a significantly increased performance.

### 3.5 Transitive conversion

Although **BioConvert** can already perform 100 conversions with 50 different formats, it also includes an experimental feature called transitive conversion that enables users to convert a file from format A to format C through an indirect route, provided that a path such as A *→ B→ C* exists. For example, if a user needs to convert a FastQ file to FAA (amino acid sequence), a user can first convert the FastQ file to a FastA file, and then use the FastA to FAA conversion. Users can also use the transitive conversion mode with the “-a” option. It should be noted, however, that information may be lost in the process (e.g., quality information is lost when converting from FastQ to FastA). Nonetheless, this this feature can be useful in certain situations.

### 3.6 A website deployment

**BioConvert** is a Python-based tool that can be installed on any modern platform with Python 3. It requires several external tools, which can be installed using Conda, for example. It may be difficult for some users to install all dependencies. Therefore, in addition to a local installation, a web-based instance of **BioConvert** is also available, hosted on the Institut Pasteur website at https://bioconvert.pasteur.cloud. The backend of the website is powered by the flask microframework to handle the upload of user’s files or dynamically proposed conversion based on the input file extension. The website is scalable and managed by Kubernetes, a container orchestration platform that automates deployment, scaling, and management of containerized applications. Nonetheless, the website uses the default conversion method only, and does not include conversions that require extra arguments. Additionally, the input file size is limited to 1GB. Therefore, for larger files or more complex computations, it is recommended to use a local installation. The website’s code source is available for customised deployment (see Data and Software Availability section).

## 4 Data and Software availability

The source code for **BioConvert** is available on GitHub under the github.com/bioconvert/bioconvert repository, with package releases posted on pypi.org website. Additionally, pre-compiled binaries are built in the Bioconda [7] project to ensure reproducibility, and Bioconda releases also provide biocontainers available on quay.io/repository/biocontainers/bioconvert website.

The source code for the **BioConvert** website can be found on https://gitlab.pasteur.fr/salsa/bioconvert.

The **BioConvert** command line can easily be parallellised and executed on HPC infrastructure using Snakemake [18] or Nextflow [22], as demonstrated in the Supplementary section.

Appropriate Apptainer containers, containing all third-party tools required by **BioConvert**, can be found on Zenodo as part of the Damona project. The **BioConvert** container DOI is https://zenodo.org/record/7704649. Additionally, the Sequana [6] project also provide a parallelized version called sequana_bioconvert, which is also available on pypi.org, and provides a simple user interface to **BioConvert**. Indeed sequana_bioconvert downloads a ready-to-use Apptainer container from the Damona project mentionned above. See Supplementary for installation and example.

## 5 Conclusion

We developed **BioConvert** in response to the dramatic increase in the number of bioinformatic formats and the growing need for multiple conversions between them. **BioConvert** facilitates transparent conversion between many formats by integrating external tools and libraries, or implementing its own simple conversions. With a consistent syntax, **BioConvert** can handle 50 file formats and 100 conversions through a command line interface. It offers both an “implicit” mode, which infers the desired conversion based on file extensions, and an “explicit” mode, in which the user specifies the conversion by name (e.g. `bam2sam`). The library also includes a benchmarking framework to determine the most efficient method for a given conversion. In addition, **BioConvert** offers a web interface that can be deployed on web servers. **BioConvert** currently covers domains such as sequencing, read alignment, variant calling, and phylogenetics, **BioConvert** is designed to be extensible and open to the integration of additional formats and conversions (e.g., proteomics, systems biology).

## Acknowledgments

HC, TC, DD and EK work has been supported by the France Génomique Consortium (ANR 10-INBS-09-08) and IBISA and the Biomics Platform of Institut Pasteur, Paris, France.

## Author contributions

HC implemented the multi benchmark, the web server, setup the CI integration and all tests for version 1.0 of **BioConvert**. TC proposed the idea of **BioConvert**, put the documentation, testing and proof of concepts. DD contributed to the web server. FL implemented the wrappers related to phylogeny, BL, YD and BB contributed to conversions and **BioConvert** core. SD refactored the core of **BioConvert** (user interface) and added conversions. All other contributors added conversions, and all authors participated in writing the manuscript.

## 6 supplementary

### 6.1 Workflows and parellelisation

Life sciences data, particularly in NGS-related fields, can be quite large, enecessitating parallelization of conversion most of the time. **BioConvert**, being a standalone application, can easily be integrated into any frmework that utilizes embarrassingly parallel paradigm.

#### 6.1.1 Snakemake example

As an illustration, we provide an example that demonstrates the usage of the Snakemake [18] framework. In this context, it is necessary to specify the input and output files, along with their respective extensions, and the conversion command.

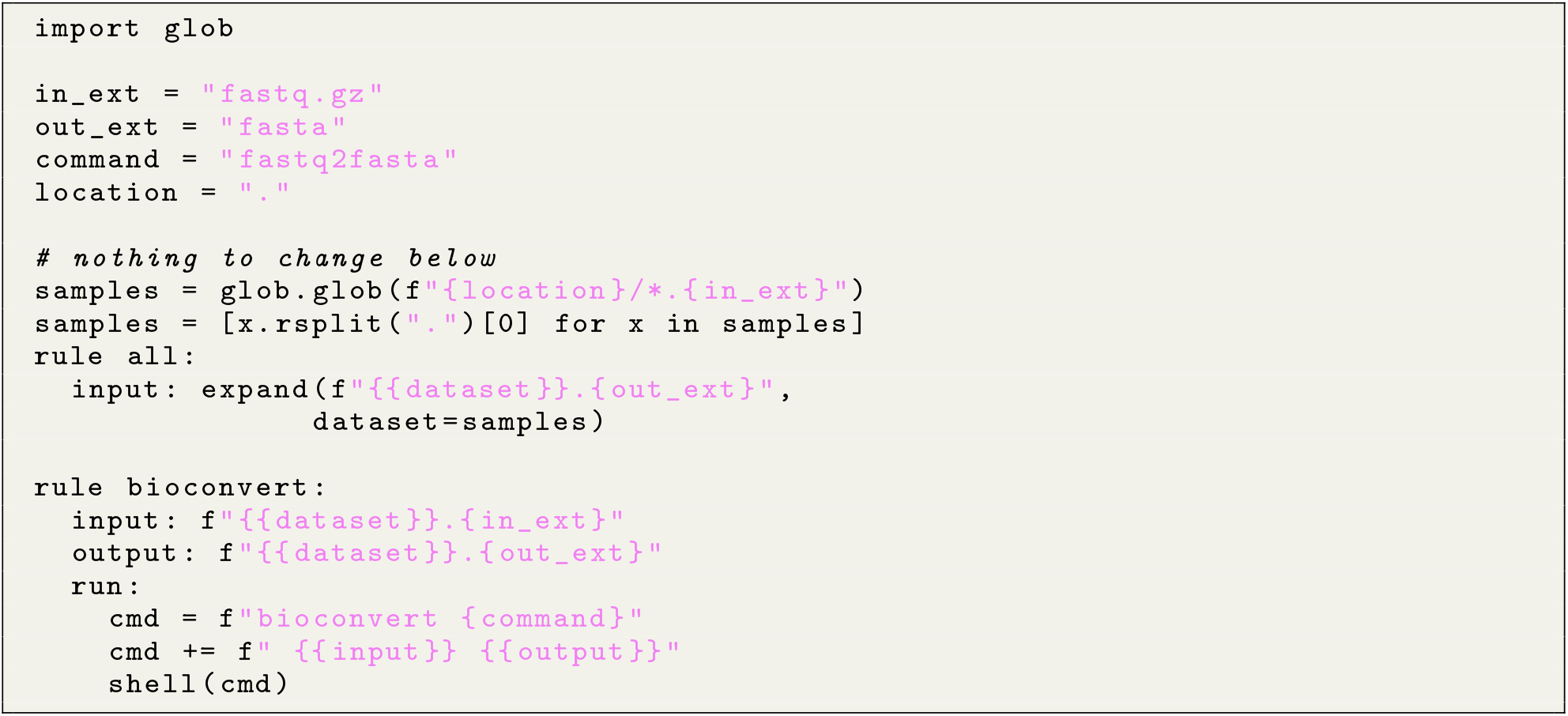

#### 6.1.2 Nextflow bioconvert pipeline

Similarly, we provide a simple example for the NextFlow [22] framework.

**Listing 8:**
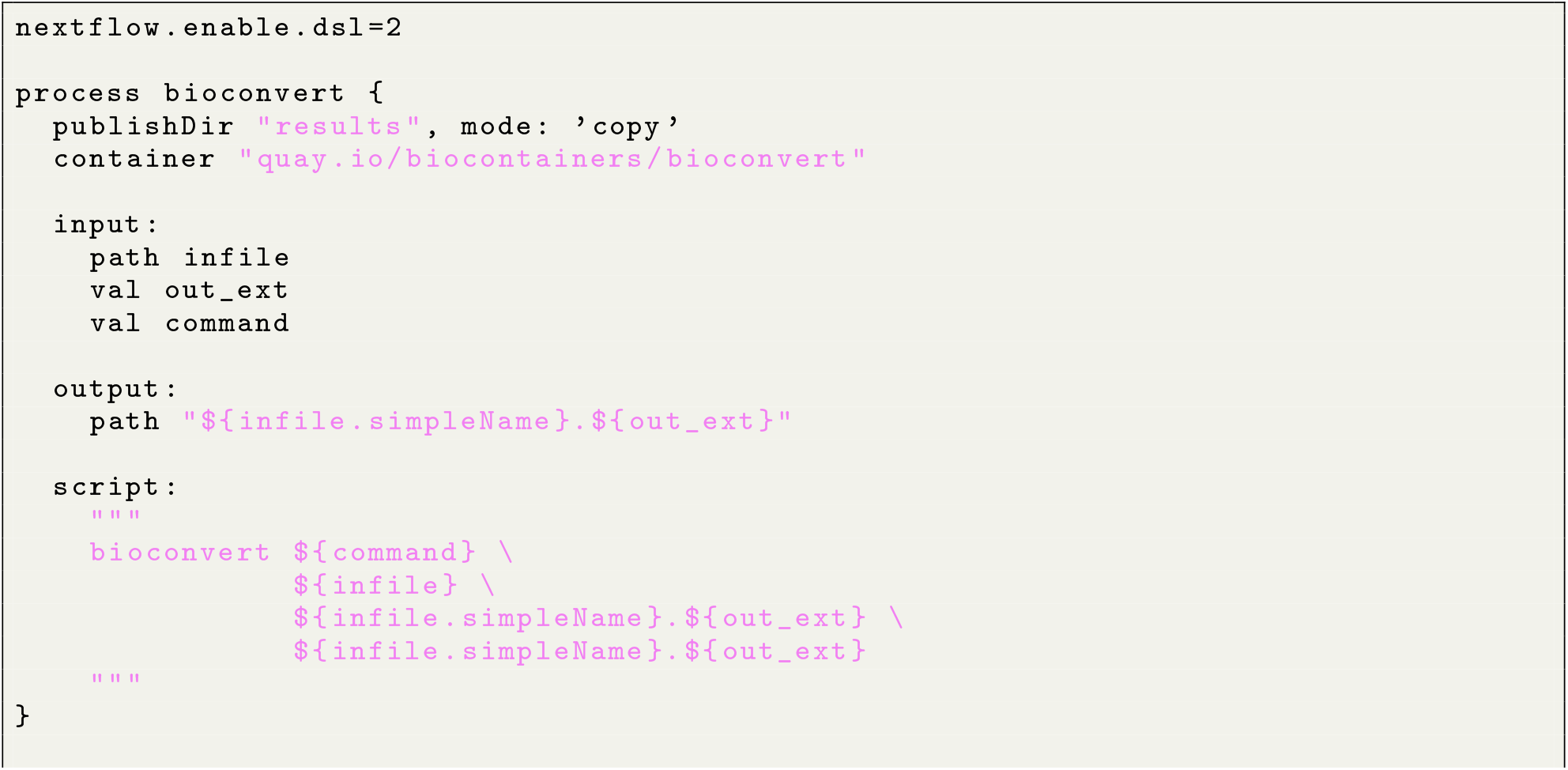
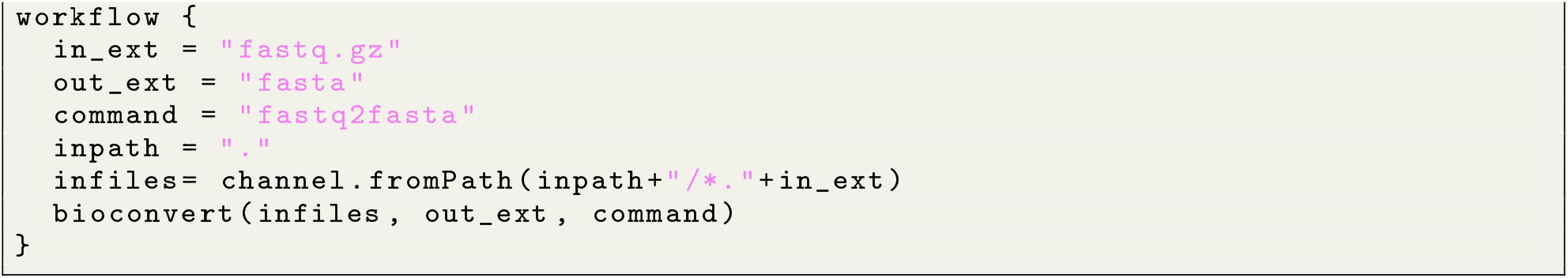
example of nextflow integration

#### 6.1.3 Sequana bioconvert pipeline

Sequana provides NGS pipelines developed in Snakemake (sequana.readthedocs.io) [6]. It also provide a pipeline dedicated to **BioConvert** that can be installed as follows:

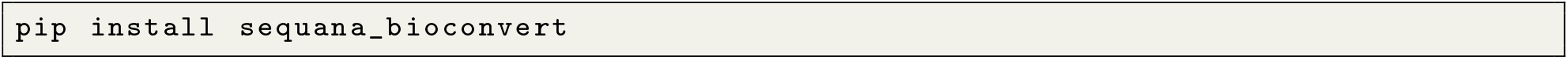

Then, a user that wishes to convert a set of BAM files into SAM files would need to type the following commands without the need to install third-party tools (here samtools) or wonder about Snakemake syntax.

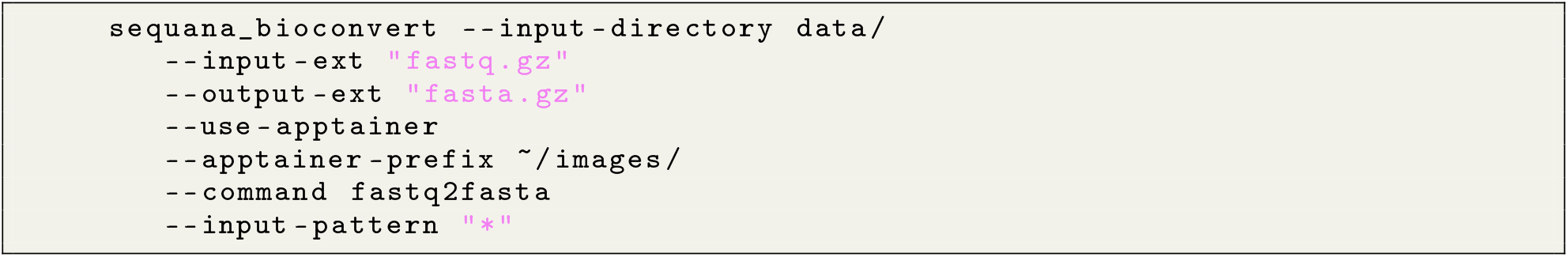

This command downloads the required container automatically and perform the conversion on the input files.

## References

[1] Lincoln Stein. Creating a bioinformatics nation. Nature, 417(6885):119–120, 2002.

[2] Simon Andrews et al. Fastqc: a quality control tool for high throughput sequence data, 2010.

[3] Heng Li, Bob Handsaker, Alec Wysoker, Tim Fennell, Jue Ruan, Nils Homer, Gabor Marth, Goncalo Abecasis, and Richard Durbin. The sequence alignment/map format and samtools. Bioinformatics, 25(16):2078–2079, 2009.

[4] Enis Afgan, Anton Nekrutenko, Bjórn A Grüning, Daniel Blankenberg, Jeremy Goecks, Michael C Schatz, Alexander E Ostrovsky, Alexandru Mahmoud, Andrew J Lonie, Anna Syme, Anne Fouilloux, Anthony Bretaudeau, Anton Nekrutenko, Anup Kumar, Arthur C Eschenlauer, Assunta D DeSanto, Aysam Guerler, Beatriz Serrano-Solano, Bérénice Batut, Björn A Grüning, Bradley W Langhorst, Bridget Carr, Bryan A Raubenolt, Cameron J Hyde, Catherine J Bromhead, Christopher B Barnett, Coline Royaux, Cristóbal Gallardo, Daniel Blankenberg, Daniel J Fornika, Dannon Baker, Dave Bouvier, Dave Clements, David A de Lima Morais, David Lopez Tabernero, Delphine Lariviere, Engy Nasr, Enis Afgan, Federico Zambelli, Florian Heyl, Fotis Psomopoulos, Frederik Coppens, Gareth R Price, Gianmauro Cuccuru, Gildas Le Corguillé, Greg Von Kuster, Gul-sum Gudukbay Akbulut, Helena Rasche, Hans-Rudolf Hotz, Ignacio Eguinoa, Igor Makunin, Isuru J Ranawaka, James P Taylor, Jayadev Joshi, Jennifer Hillman-Jackson, Jeremy Goecks, John M Chilton, Kaivan Kamali, Keith Suderman, Krzysztof Poterlowicz, Le Bras Yvan, Lucille Lopez-Delisle, Luke Sargent, Madeline E Bassetti, Marco Antonio Tangaro, Marius van den Beek, Martin Cêch, Matthias Bernt, Matthias Fahrner, Mehmet Tekman, Melanie C Föll, Michael C Schatz, Michael R Crusoe, Miguel Roncoroni, Natalie Kucher, Nate Coraor, Nicholas Stoler, Nick Rhodes, Nicola Soranzo, Niko Pinter, Nuwan A Goonasekera, Pablo A Moreno, Pavankumar Videm, Petera Melanie, Pietro Mandreoli, Pratik D Jagtap, Qiang Gu, Ralf J M Weber, Ross Lazarus, Ruben H P Vorderman, Saskia Hiltemann, Sergey Golitsynskiy, Shilpa Garg, Simon A Bray, Simon L Gladman, Simone Leo, Subina P Mehta, Timothy J Griffin, Vahid Jalili, Vandenbrouck Yves, Victor Wen, Vijay K Nagampalli, Wendi A Bacon, Willem de Koning, Wolfgang Maier, and Peter J Briggs. The galaxy platform for accessible, reproducible and collaborative biomedical analyses: 2022 update. Nucleic Acids Research, 50(W1):W345–W351, April 2022.

[5] Philip A Ewels, Alexander Peltzer, Sven Fillinger, Harshil Patel, Johannes Alneberg, Andreas Wilm, Maxime Ulysse Garcia, Paolo Di Tommaso, and Sven Nahnsen. The nf-core framework for community-curated bioinformatics pipelines. Nature biotechnology, 38(3):276–278, 2020.

[6] Thomas Cokelaer, Dimitri Desvillechabrol, Rachel Legendre, and Mélissa Cardon. ‘sequana’: a set of snakemake ngs pipelines. Journal of Open Source Software, 2(16):352, 2017.

[7] Björn Grüning, Ryan Dale, Andreas Sjödin, Brad A Chapman, Jillian Rowe, Christopher H Tomkins-Tinch, Renan Valieris, Johannes Köster, and Bioconda Team. Bioconda: sustainable and comprehensive software distribution for the life sciences. Nature methods, 15(7):475–476, 2018.

[8] Jon Ison, Matúš Kalaš, Inge Jonassen, Dan Bolser, Mahmut Uludag, Hamish McWilliam, James Malone, Rodrigo Lopez, Steve Pettifer, and Peter Rice. Edam: an ontology of bioinformatics operations, types of data and identifiers, topics and formats. Bioinformatics, 29(10):1325–1332, 2013.

[9] Jon Ison, Hans Ienasescu, Piotr Chmura, Emil Rydza, Hervé Ménager, Matúš Kalaš, Veit Schwämmle, Björn Grüning, Niall Beard, Rodrigo Lopez, et al. The bio. tools registry of software tools and data resources for the life sciences. Genome biology, 20(1):1–4, 2019.

[10] Fábio Madeira, Matt Pearce, Adrian R N Tivey, Prasad Basutkar, Joon Lee, Ossama Edbali, Nandana Madhusoodanan, Anton Kolesnikov, and Rodrigo Lopez. Search and sequence analysis tools services from embl-ebi in 2022. Nucleic acids research, page gkac240, April 2022.

[11] Thomas Cokelaer, Dennis Pultz, Lea M Harder, Jordi Serra-Musach, and Julio Saez-Rodriguez. Bioservices: a common python package to access biological web services programmatically. Bioinformatics, 29(24):3241– 3242, 2013.

[12] Nikhil Paul, Markus W Schmitt, and Karl Holzmann. Nanopore sequencing: Principles, applications, and challenges. Frontiers in Genetics, 11:612, 2020.

[13] Helle Madsen, Ralf Kaiser, Anna Sagyan, Rasmus Sørensen, Thomas F Dyrlund, Henrik Lund, Martin R Larsen, Markus Taubert, and Peter R Nielsen. Pacbio sequencing using the smrt technology. Methods, 59(1):1–11, 2013.

[14] Don Gilbert. Sequence file format conversion with command-line readseq. Current protocols in bioinformatics, (1):A–1E, 2003.

[15] Nicolas Rodriguez, Jean-Baptiste Pettit, Piero Dalle Pezze, Lu Li, Arnaud Henry, Martijn P van Iersel, Gael Jalowicki, Martina Kutmon, Kedar N Natarajan, David Tolnay, et al. The systems biology format converter. BMC bioinformatics, 17(1):1–7, 2016.

[16] Frédéric Lemoine and Olivier Gascuel. Gotree/goalign: toolkit and go api to facilitate the development of phylogenetic workflows. NAR Genomics and Bioinformatics, 3(3):lqab075, 2021.

[17] Derek Draper and James Smith. Bamtools: a c++ api and toolkit for reading, writing, and manipulating bam files. Bioinformatics, 27(5):778–779, 2011.

[18] F Mölder, KP Jablonski, B Letcher, MB Hall, CH Tomkins-Tinch, V Sochat, J Forster, S Lee, SO Twardziok, A Kanitz, A Wilm, M Holtgrewe, S Rahmann, S Nahnsen, and J Köster. Sustainable data analysis with snake-make. F1000Research, 10(33), 2021.

[19] Nicholas J Loman, Joshua Quick, Jared T Simpson, Clint Durrant, Mitchell Loose, James Stalker, Thomas R Connor, David R Bentley, and Sarah R Harris. Mosdepth: Fast computation of read depth for wgs, exome and target capture datasets. Bioinformatics, 33(16):2556–2558, 2017.

[20] Heng Li. seqtk toolkit for processing sequences in fasta/q formats, 2012.

[21] Wei Shen, Shuai Le, Yan Li, and Fuquan Hu. Seqkit: a cross-platform and ultrafast toolkit for fasta/q file manipulation. PloS one, 11(10):e0163962, 2016.

[22] P. et al. Di Tommaso. Nextflow enables reproducible computational workflows. Nature Biotechnology, 35, 2017.

